# Investigating the relationship between phoneme categorization and reading ability

**DOI:** 10.1101/305748

**Authors:** Gabrielle E. O’Brien, Daniel R. McCloy, Emily C. Kubota, Jason D. Yeatman

**Affiliations:** Institute for Learning and Brain Sciences and Department of Speech and Hearing Sciences, University of Washington, Seattle, WA, USA

## Abstract

Dyslexia is associated with abnormal performance on many auditory psychophysics tasks, particularly those involving the categorization of speech sounds. However, it is debated whether those apparent auditory deficits arise from (a) reduced sensitivity to particular acoustic cues, (b) the difficulty of experimental tasks, or (c) unmodeled lapses of attention. Here we investigate the relationship between phoneme categorization and reading ability, with special attention to the nature of the cue encoding the phoneme contrast (static versus dynamic), differences in task paradigm difficulty, and methodological details of psychometric model fitting. We find a robust relationship between reading ability and categorization performance, show that task difficulty cannot fully explain that relationship, and provide evidence that the deficit is not restricted to dynamic cue contrasts, contrary to prior reports. Finally, we demonstrate that improved modeling of behavioral responses suggests that the gap between dyslexics and typical readers may be smaller than previously reported.

## Introduction

Dyslexia is a learning disability that affects between 5% and 17% of the population and poses a substantial economic and psychological burden for those affected (Shaywitz, 1998; Snowling, 2000; Lyon, Shaywitz and Shaywitz, 2003). Despite decades of research, it remains unclear why so many children without obvious intellectual, sensory, or circumstantial challenges find written word recognition so difficult.

One popular and persistent theory is that dyslexia arises as a result of an underlying auditory processing deficit (Tallal, 1980; Farmer and Klein, 1995; Snowling, 1998; Van Ingelghem, Mieke, Boets *et al.*, 2005; Goswami, 2011; Steinbrink *et al.*, 2014). According to this theory, a low-level auditory processing deficit disrupts the formation of a child’s internal model of speech sounds (phonemes) during early language learning; later, when young learners attempt to associate written letters (graphemes) with phonemes, they struggle because their internal representation of phonemes is compromised (Poelmans *et al.*, 2011).

In line with this hypothesis, many studies do report group differences between dyslexics and typical reading control participants in auditory psychophysical tasks including amplitude modulation detection (McAnally and Stein, 1997; Witton, 1998; Menell, McAnally and Stein, 1999; Rocheron *et al.*, 2002; Hämäläinen *et al.*, 2009), frequency modulation detection (Witton *et al.*, 2002; Gibson, Hogben and Fletcher, 2006; Stoodley *et al.*, 2006; Boets *et al.*, 2007; Dawes *et al.*, 2009), rise time discrimination, and duration discrimination (Banai and Ahissar, 2004, 2006; Thomson *et al.*, 2006; Thomson and Goswami, 2008; Goswami *et al.*, 2011). Moreover, attributing dyslexia to an auditory deficit is appealing because one of the most effective predictors of reading difficulty is poor phonemic awareness—the ability to identify, segment and manipulate the phonemes within a spoken word (Bus and van IJzendoorn, 1999; Hulme *et al.*, 2002). If a child’s phoneme representation were abnormal due to auditory processing deficits, it could be a common factor underlying both poor performance in auditory phoneme awareness tests and difficulties learning to decode written words among children with dyslexia.

In one variant of this auditory hypothesis, dyslexia is thought to involve a deficit specifically in the processing of rapid modulations in sound, usually referred to as a “rapid temporal processing” deficit (Tallal, 1980; Merzenich *et al.*, 1996; Tallal *et al.*, 1996). This idea has been controversial (see Rosen, 1999, 2003 for review), but also remains as one of the widely cited accounts of auditory deficits in dyslexia (Van Ingelghem, Mieke, Boets *et al.*, 2005). One source of controversy is that the rapid temporal processing deficit is far from universal; in Tallal’s (1980) study that first proposed a causal relationship between rapid temporal processing and reading ability, only 8 out of 20 dyslexic children showed a deficit on a temporal order judgment task. Similarly, in a more comprehensive study of 17 dyslexic adults, only 11 had impaired performance on an extensive battery of auditory psychophysical tasks (Ramus *et al.*, 2003). Moreover, in that study, tasks requiring temporal cue sensitivity were not systematically more effective than non-temporal tasks at separating dyslexics from controls. More recently, the rapid temporal processing hypothesis has been reframed as a deficit in processing dynamic, but not necessarily rapid, aspects of speech (Boets *et al.*, 2011; Poelmans *et al.*, 2011; Law *et al.*, 2014).

On the whole, there is ample evidence that abnormal performance on auditory psychophysical tasks is frequently associated with dyslexia (Hämäläinen, Salminen and Leppänen, 2013), but much less evidence that a specific deficit in temporal processing is causally related to dyslexia (or even consistently co-occurring with it). But regardless of whether the alleged deficit involves all modulations of the speech signal or only rapid ones, there are at least two major reasons why the theory that auditory temporal processing deficits cause the reading difficulties seen in dyslexia has been called into question. First, although correlations have been found between certain psychoacoustic measures and reading skills (Witton, 1998; Goswami *et al.*, 2002; Vandermosten *et al.*, 2010), there have been no direct findings relating performance on psychoacoustic tasks to phonological representations (Rosen, 2003). On the contrary, there is growing evidence that poor performance on psychophysical tasks may be partially or totally explained by the demands of the test conditions and not the stimuli themselves (Banai and Ahissar, 2006). In this view, it is not fuzzy phonological representations, but rather poor working memory—which often co-occurs with dyslexia (Siegel and Ryan, 1989; Swanson, 1993; Vargo, Grosser and Spafford, 1995; Wang and Gathercole, 2013)—that drives deficits in both psychoacoustic task performance and in reading ability (Amitay *et al.*, 2002; Banai and Ahissar, 2004). Proponents of this hypothesis argue that the phonological deficit associated with dyslexia is only observed when considerable memory and time constraints are imposed by the task (Ramus and Szenkovits, 2008); evidence for this view comes from studies in which listeners perform different kinds of discrimination paradigms on the same auditory stimuli (Ahissar *et al.*, 2006; Banai and Ahissar, 2006; Ahissar, 2007; Ziegler, 2008). Dyslexia is also known to be comorbid with ADHD, and numerous reviews have suggested that, in addition to working memory, differences in attention may be another key driver of the observed group differences (Ramus *et al.*, 2003; Roach, Edwards and Hogben, 2004).

A second reason to question the link between auditory processing deficits and dyslexia is that the standard technique for data analysis in psychophysical experiments is prone to severe bias. Specifically, in many phoneme categorization studies, participants identify auditory stimuli that vary along some continuum as belonging to one of two groups, a psychometric function is fit to the resulting data, and the slopes of these psychometric functions are compared between groups of dyslexic and control readers (for reviews, see Vandermosten *et al.*, 2011; Noordenbos and Serniclaes, 2015). In this approach, a steep slope indicates a clear boundary between phoneme categories, whereas a shallow slope suggests less defined categories (i.e., fuzzy phonological representations), possibly due to poor sensitivity to the auditory cue(s) that mark the phonemic contrast. Unfortunately, most studies in the dyslexia literature fit the psychometric function using algorithms that fix its asymptotes at zero and on, which is equivalent to assuming that categorization performance is perfect at the extremes of the stimulus continuum (i.e., assuming a “lapse rate” of zero). This assumption is questionable in light of the evidence that attention, working memory, or task difficulty, rather than stimulus properties, may underlie group differences between readers with dyslexia and control subjects.

The zero lapse rate assumption is particularly problematic given that fixed asymptotes at zero and one leads to strongly downward-biased slope estimates even when true lapse rates are fairly small (Wichmann and Hill, 2001a, 2001b). In other words, if a participant makes chance errors on categorization, and these errors happen on trials near the ends of the continuum, the canonical psychometric fitting routine used in most previous studies will underestimate the steepness of the category boundary. Thus, a tendency to make a larger number of random errors (due to inattention, memory, or task-related factors) will be wrongly attributed to an indistinct category boundary on the contrast under study. Since it is precisely the subjects with dyslexia that more frequently show attention or working memory deficits, research purporting to show less distinct phoneme boundaries in readers with dyslexia may in fact reflect non-auditory, non-linguistic differences between study populations.

Unfortunately, most studies in the dyslexia literature that use psychometric functions to model categorization performance appear to suffer from this bias. Although most do not report their analysis methods in sufficient detail to be certain, we infer (based on published plots, and the lack of any mention of asymptote estimation) that the bias is widespread (e.g., Reed, 1989; Manis *et al.*, 1997; Breier *et al.*, 2001; Chiappe, Chiappe and Siegel, 2001; Maassen *et al.*, 2001; Bogliotti *et al.*, 2008; Zhang *et al.*, 2012). A few studies report using software that in principle supports asymptote estimation during psychometric fitting, but the authors did not report the parameters used in their analysis (e.g., Vandermosten *et al.*, 2010, 2011). In one case, researchers who re-fit their psychometric curves without data from the continuum endpoints found a reduced effect size, prompting the authors to wonder whether any effect would remain if an unbiased estimator of slope were used (Messaoud-Galusi, Hazan and Rosen, 2011).

In light of all these problems—the inhomogeneity of findings, confounding influences of experimental task design, and bias introduced by analysis choices—it is reasonable to wonder whether children with dyslexia really have any abnormality in phoneme categorization, once those factors are all controlled for. The present study addresses the relationship between reading ability and phoneme categorization ability, and in particular, whether children with dyslexia show a greater deficit on phoneme contrasts that rely on temporally varying (dynamic) cues, as opposed to static cues. This study avoids the aforementioned methodological problems by using multiple paradigms with different attentional and memory demands, and by analyzing categorization performance using Bayesian estimation of the psychometric function (Schütt *et al.*, 2015). In this approach, the four parameters of a psychometric function—the threshold, slope, and two asymptote parameters—are assigned prior probability distributions, which formalize the experimenter’s assumptions about their likely values. By allowing the asymptote parameters to vary, the slope is estimated in a less-biased way than in traditional modeling approaches with fixed asymptotes. However, because fitting routines trade off between asymptote parameters and the slope parameter of a logistic model in optimization space (Treutwein and Strasburger, 1999), it can be difficult to estimate both accurately at the same time. To address this difficulty, we first performed cross-validation on the prior distribution of the asymptote parameters to determine the optimal model to fit the data.

This paper presents data from 44 children, aged 8–12 years. Our experimental task is based on the design of Vandermosten et al. (2010, 2011), which assessed categorization performance for two kinds of stimulus continua: those that differed based on a spectrotemporal cue (dynamic), and those that differed based on a purely spectral cue (static). In the original study, the authors concluded that children with dyslexia are specifically impaired at categorizing sounds (both speech and non-speech) that differ on the basis of dynamic cues. However, although the dynamic and static stimuli in their study were equated for overall length, the duration of the cues relevant for categorization were not equal: in the dynamic stimuli, the cue (a vowel formant transition) was available for 100 ms, but in the static stimuli, the cue (the formant frequency of a steady state vowel) was available for 350 ms. This raises the question of whether cue duration, rather than the dynamic nature of the cue, was the source of apparent impairment in categorization among participants with dyslexia. The present study avoids this confound by changing the “static cue” stimuli from steady-state vowels to fricative consonants (a /∫a/~/sa/ continuum), so that the relevant cue duration is 100 ms in both the static (/∫a/~/sa/) and dynamic (/ba/~/da/) stimulus continua. Additionally, Vandermosten and colleagues used a test paradigm in which listeners heard three sounds and were asked to decide if the third sound was more like the first or second (an ABX design). Here we included both an ABX task and a single-stimulus categorization task, to see whether the memory and attention demands of the ABX paradigm may have played a role in previous findings. Thus, by (a) assessing categorical perception of speech continua with static and dynamic cues, (b) varying the cognitive demands of the psychophysical paradigm, and (c) empirically determining the optimal parameterization of the psychometric function with cross-validation, we aim to clarify the role of auditory processing deficits in dyslexia. Note that for visualization purposes we have divided the participants into three groups (Dyslexic, Below Average, and Above Average), but all of our statistical analyses treat reading ability as a continuous variable, represented by a composite reading score (see Materials & Methods for details).

## Results

Response functions for each subject on the two stimulus continua (aggregated across task paradigms) are shown in Figure 1. For Above Average readers, response functions show the typical sigmoid shape, with a steep boundary between categories and consistent labeling within categories; for readers in the Below Average and Dyslexic groups, response functions are much more heterogeneous, with some subjects exhibiting inconsistent performance even at continuum endpoints. The group means are suggestive of differences in both psychometric slope and lapse rate on both the static (/∫a/~/sa/) and dynamic (/ba/~/da/) cue continua.

**Figure 1.**
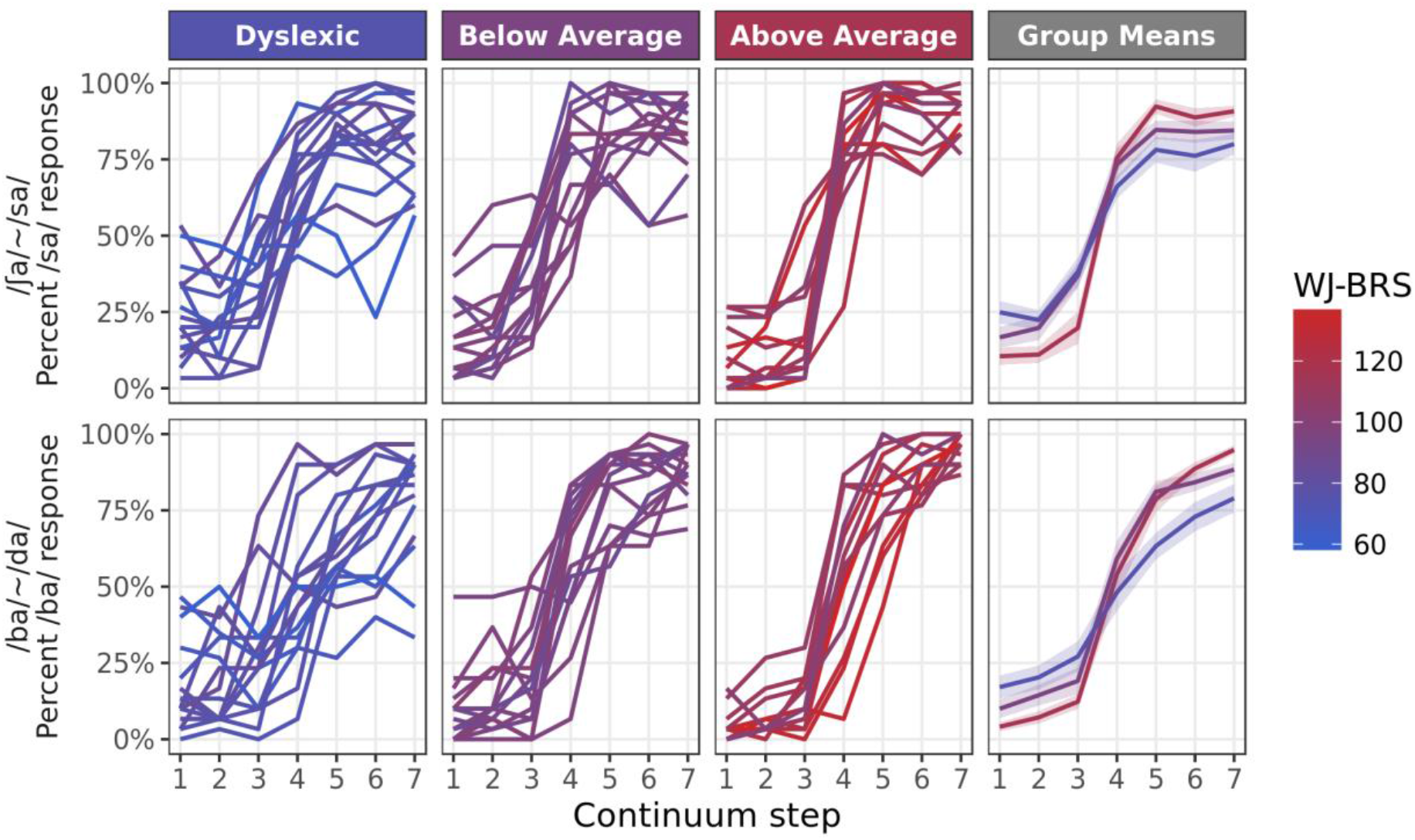
*Response functions for each subject and each speech continuum. Results are averaged over responses from both the ABX and single-interval paradigms. The color of each line maps to the subject’s reading ability based on the Woodcock Johnson Basic Reading Skills Composite score (WJ-BRS)*.

Figure 2 shows the psychometric functions fit to each subject’s response data on the two continua (aggregated across paradigms). Unsurprisingly, the fitted curves reflect the same pattern seen in the raw data: subjects with high reading scores tend to show consistent categorization at stimulus endpoints and steep category transitions near the center of the stimulus continuum, whereas subjects with poor reading scores show a diversity of curve shapes. For some subjects with poor reading scores the psychometric curves appear quite similar to those of the skilled readers, while others exhibit shallow slopes at the category transition and/or high lapse rates at the continuum endpoints. Estimates of the category boundary (the “threshold” parameter of the psychometric) also appear to span a wider range of continuum values among poor readers compared with readers in the above average group.

**Figure 2.**
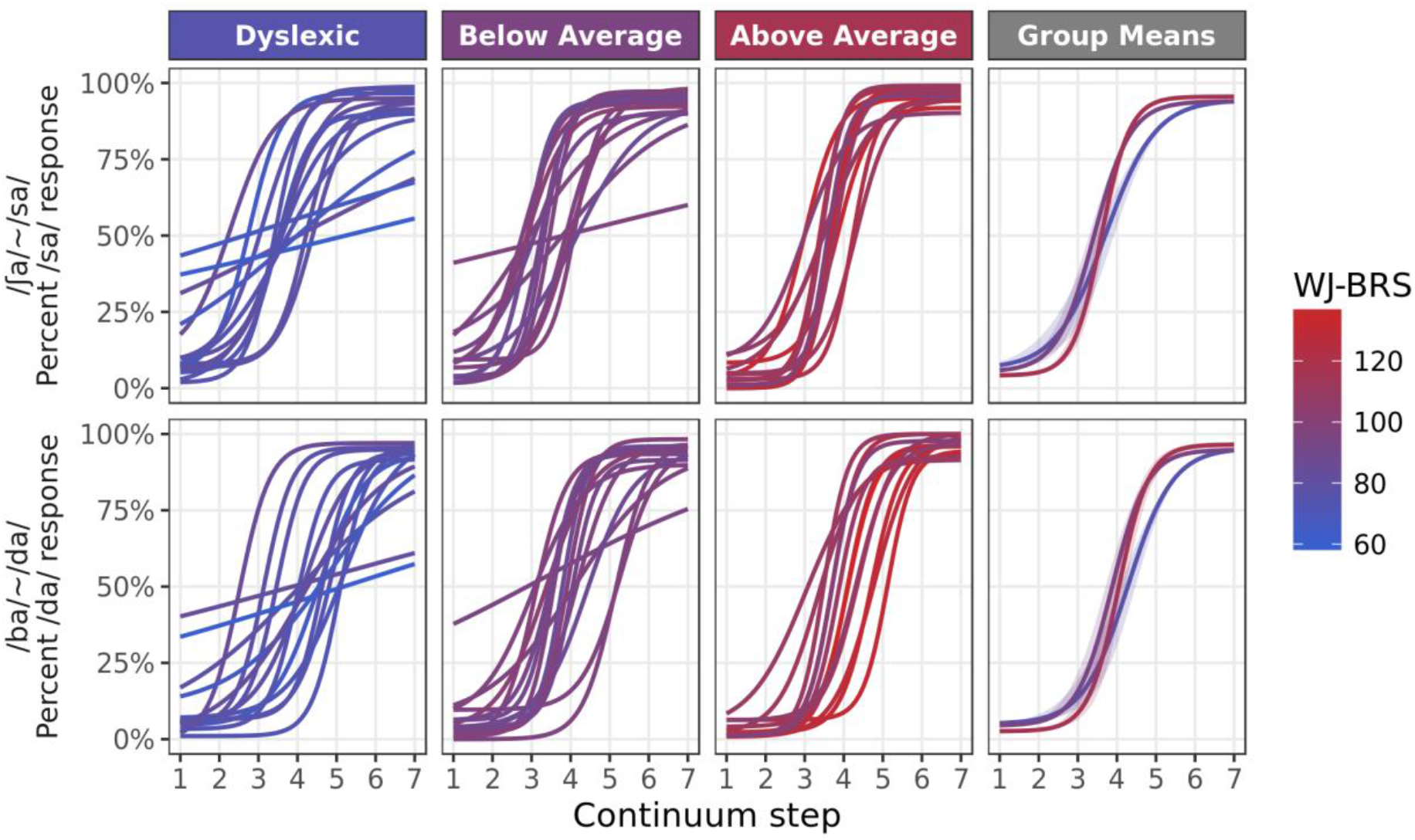
*Fitted response functions for each subject and each continuum. Parameters (slope, threshold, and the two asymptotic parameters) were averaged over estimates from both the ABX and single-interval paradigms. The color of each line maps to the subject’s reading ability (WJ-BRS score)*.

### High reading score predicts reliable phoneme categorization

The group-level patterns of slope and lapse rate estimates are summarized in Figure 3. There is a clear trend of increasingly steeper slopes for children with higher reading scores, across both stimulus continua and both experimental paradigms. The pattern of lapse rate estimates is less consistent, though the lapse rate estimates tended to be higher for poor readers. Slope and lapse rate estimates were each modeled using mixed effects regression, as implemented in the lme4 library for R (Bates *et al.*, 2015). Fixed-effect predictors with deviation coding were used for both the continuum (static /∫a/~/sa/ versus dynamic /ba/~/da/) and paradigm (ABX versus single-stimulus) variables, and reading ability (WJ-BRS score) was included as a continuous fixed-effect predictor. A random intercept by participant was also included. Fixed-effect predictors were also included to control for the effects of ADHD diagnosis and non-verbal IQ (age-normalized WASI Matrix Reasoning score). Because two blocks of the ABX test were administered per subject, slope or lapse rate parameter estimates were averaged over the two blocks for each individual before regression modeling.

**Figure 3.**
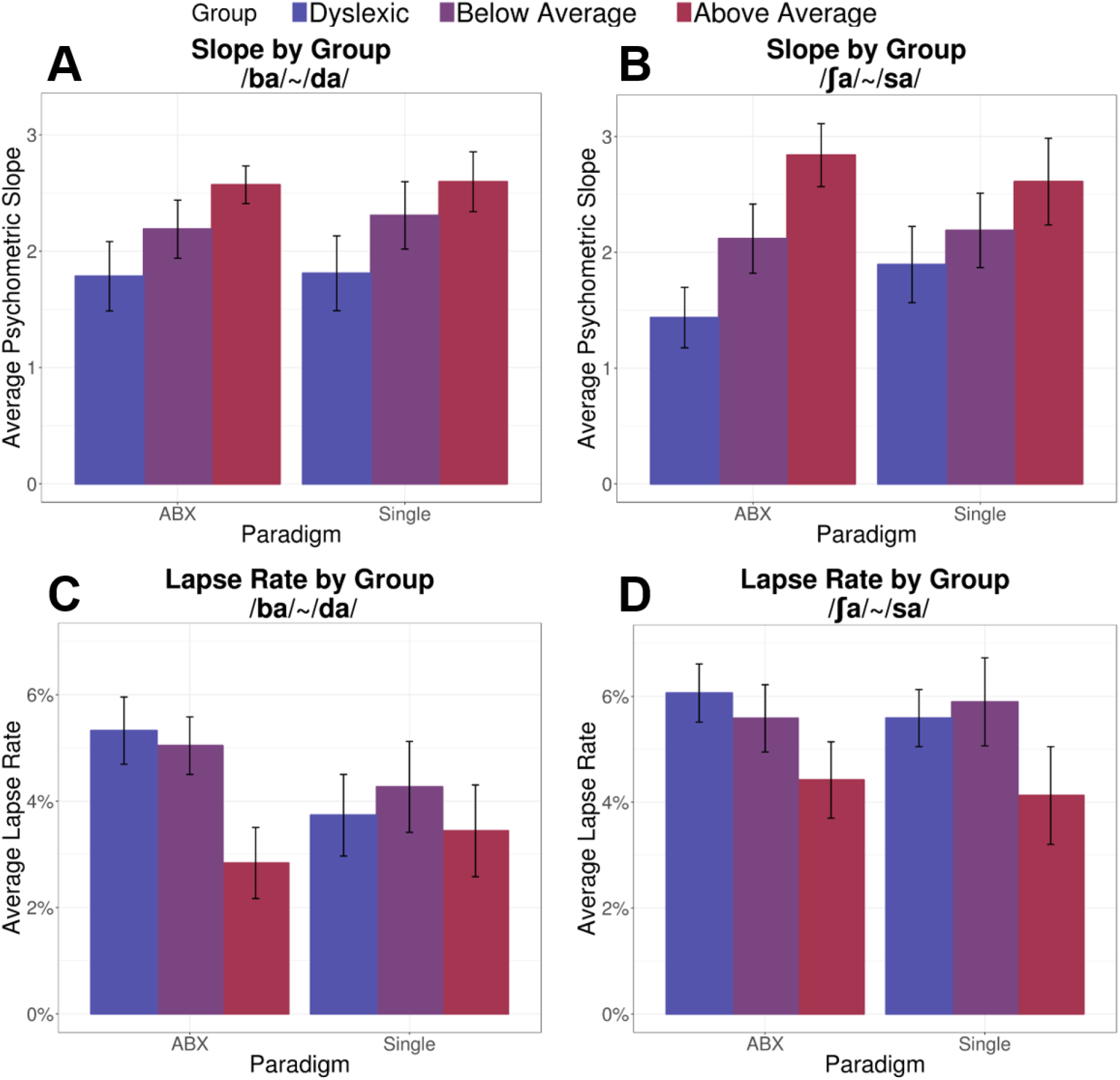
*Average psychometric slope within groups for the /ba/~/da/ and /ſa/~/sa/ continua are shown in panels A and B, separated by paradigm. Average lapse rate parameters for both continua are shown in Panels C and D. Error bars represent one standard error of the mean*.

The fully-specified model of psychometric slope estimates indicated that only one predictor—WJ-BRS reading score—had a significant effect on slope estimates; there was no reliable difference in psychometric slopes between the static and dynamic cue continua, nor was there a reliable difference between ABX blocks and single-interval blocks. Stepwise nested model comparisons were then used to eliminate irrelevant model parameters and yield the most parsimonious model (see supplement for details). The most parsimonious model of psychometric slope contained only a continuous predictor of reading ability (WJ-BRS) and a random intercept for each participant (see Table 1). The effect size of reading ability on psychometric slope was relatively modest; a 1-unit improvement in WJ-BRS score was associated with an increase of 0.023 in the slope of the psychometric function indicating that poor readers tend to have shallower psychometric slopes (p<0.001).

To model the relationship between lapse rate and reading score, we first combined the two asymptote estimates from the psychometric fits, by averaging the asymptote deviation from zero or one (i.e., a lower asymptote of 0.1 and an upper asymptote of 0.9 both correspond to a lapse rate of 0.1; the upper and lower lapse rates were averaged for each psychometric fit). The same initial model specification and simplification procedure used with the slope model was also used for the lapse rate model. The final model of lapse rate contained a continuous predictor for reading ability, a categorical predictor for continuum, and a random effect for participant (see Table 1); no reliable effect of task paradigm was detected. Higher reading ability was associated with slightly smaller lapse rates (about 0.04% smaller per 1-unit increase of WJ-BRS score), and the static /∫a/~/sa/ continuum was associated with larger lapse rates (about 1% higher than the lapse rate estimates for the /ba/~/da/ continuum). Thus, compared to strong readers, poor readers show shallower psychometric slopes and are also more likely to make errors even near the continuum endpoints, and there is a general tendency for all participants to make more errors near continuum endpoints with the static-cue stimuli than with the dynamic-cue stimuli.

### Traditional model fitting misrepresents effect of cue continuum

For comparison with the methods commonly used in prior studies, we also fit a mixed effects model using slope estimates from a traditional 2-parameter psychometric function fit (with asymptotes fixed at 0 and 1), and performed the same model simplification procedure. In this case, the most parsimonious model indicated a significant effect of stimulus continuum on psychometric slope, in addition to the effect of reading ability seen in the model of 4-parameter fits (see Table 1). The magnitude of the estimated effect of continuum on psychometric slope is quite large (difference in slope of 0.167 between dynamic and static cue continua, comparable to a difference of about 10 points on the WJ-BRS), with lower slopes associated with the static-cue (/∫a/~/sa/) continuum. Comparing this to the models of 4-parameter fits, we see reasonably similar estimates of the effect of reading ability on psychometric slope, but the effect of stimulus continuum is allocated differently depending on the underlying assumptions of the psychometric fitting routine.

**Table 1.** *Summary of significant model parameters for the four models*.

| Model | Parameter | $\beta$ | SE | *p* |
| --- | --- | --- | --- | --- |
| 4-parameter slope | WJ-BRS | 0.023 | 0.006 | 0.0005 |
| 4-parameter lapse | WJ-BRS | -0.0004 | 0.0001 | 0.002 |
|  | Continuum (static) | 0.010 | 0.003 | 0.007 |
| 2-parameter slope | WJ-BRS | 0.017 | 0.004 | 0.0004 |
|  | Continuum (static) | -0.167 | 0.078 | 0.039 |
| 1<sup>st</sup> principal component | WJ-BRS | 0.021 | 0.006 | 0.0007 |
|  | WJ-BRS:Paradigm | 0.012 | 0.006 | 0.048 |

### Principal component analysis reveals an effect of task paradigm

Although slope and lapse rate can be thought of as representing separate cognitive processes (category discrimination and attention, respectively), in reality it is difficult to disentangle what is being modeled by each parameter: the two are not truly independent in the psychometric fitting procedure. Moreover, in speech continua, slope and lapse rate may be influenced by the same underlying processes: the endpoints are determined by natural speech tokens, which may not be reliably categorized 100% of the time by all subjects, due to the complex relationships among speech cues, talker/listener dialect, context, etc. For these reasons, we also performed a principal components analysis that combined slope and asymptote estimates into a single index of psychometric curve shape. The first principal component was a linear combination of the slope and both asymptotes (slope: 0.678; lower asymptote: −0.268; upper asymptote: −0.685) that explains about 41% of variance in the data. Using an identical modeling procedure to the analysis of slope and lapse rate, we iteratively eliminated predictors from a fully specified mixed model of the first principal component. The most parsimonious model showed a main effect of reading ability (WJ-BRS score) and a significant interaction between reading ability and experimental paradigm, but no effect of stimulus continuum (see Table 1). The sign of the interaction indicated a greater difference of curve shape between task paradigms in poor readers than in strong readers. Thus, in line with previous hypotheses (Banai and Ahissar, 2006; Ramus and Szenkovits, 2008), this model suggests that the specific paradigm used in a psychophysical experiment can amplify deficits in people with dyslexia. However, dyslexics still perform poorly on phoneme categorization tasks irrespective of the paradigm used in the experiment.

### Poor readers struggle even with unambiguous stimuli at continuum endpoints

The lapse rate results discussed above suggest that children with dyslexia may have considerable difficulty even in categorizing unambiguous stimuli at the continuum endpoints. We performed post-hoc analyses of accuracy and reaction time to further investigate this possibility. In particular, we were interested in whether the high lapse rate estimates resulted from poor readers’ performance degrading over the course of an experimental block. We were also interested in whether the pattern of reaction times across different parts of the stimulus continuum varied systematically with reading ability.

#### Accuracy at continuum endpoints

Because our young subjects ranged considerably in attention span and motivation, we analyzed the stationarity of response accuracy at continuum endpoints, to ascertain whether task performance declined from start to finish. Following the design of Messaoud-Galusi et al (2011), for each test block (6 total per subject) we computed the number of correct assignments of the stimulus endpoints. There were twenty endpoint stimulus presentations per block.

We then modeled correct endpoint stimulus categorization as a function of trial number (1 through 70), reading score, stimulus continuum, and test paradigm (ABX or Single-stimulus), plus covariates for age and ADHD diagnosis (random effect of subject). WJ-BRS, age, and trial number were centered for clear interpretation of interaction terms. Following the same model fitting and simplification procedures described above, we found the most parsimonious model included only main effects of WJ-BRS (β=0.004, SE = 0.001, p = 0.001) and trial number (β=−0.0008, SE = 0.0003, p = 0.022). The main effect of WJ-BRS indicated that poor readers were more likely than typical readers to err when categorizing unambiguous stimuli, echoing the earlier analysis of lapse rate. The main effect of trial number indicates that all readers tended to decline in accuracy as the test block progressed. The lack of a significant interaction between trial number and WJ-BRS indicates that on average, dyslexics were no more likely than other subjects to decline in performance due to fatigue or inattention as the test block wore on. Thus, inability to stay focused over the course of a psychophysics experiment does not explain phoneme categorization deficits in children with dyslexia.

#### Reaction time analysis

With stimulus continua that span a categorical boundary, it is expected that reaction times will be longer for stimuli near the category boundary (Pisoni and Tash, 1974; Repp, 1981), reflecting the perceptual uncertainty and resulting decision-making difficulty associated with ambiguous stimuli. Although our participants were not under time constraints in this study, they were encouraged to respond as soon as they could after stimulus presentation, and reaction times were recorded. To see whether the children we tested showed the expected pattern (faster reaction times on unambiguous stimuli at or near continuum endpoints, and slower reaction times on ambiguous stimuli near continuum centers), we modeled reaction times across continuum steps with a second-order polynomial basis (and its interaction with reading ability).

Kernel density estimates of the reaction time distributions for the three groups are shown in Figure 4A, individual and group-level polynomial fits are shown in Figure 4B and C, and the quadratic coefficients of the polynomial fits are shown in Figure 4D. The figure separates data from the two stimulus continua for easier comparison with Figures 1 and 2, but since we had no specific hypothesis about reaction time differences between continua, we did not include that comparison in the post-hoc analysis.

Model results showed an overall relationship between reading ability and reaction time (β = −0.0025, SE = 0.0007, p < 0.001) indicating an average 2.5 ms speed-up in reaction time per unit change in WJ-BRS score, although there was substantial overlap in reaction time distributions across groups (cf. Figure 4A). The quadratic polynomial coefficient was also significant (β = −2.196, SE = 0.365, p < 0.001) showing a general tendency for faster responses at continuum endpoints. Finally, there was a significant interaction between the quadratic polynomial coefficient and WJ-BRS score (β = −0.066, SE = 0.019, p < 0.001), indicating a more downward-curving polynomial for subjects with higher reading scores (cf. Figure 4B, C, and D). In other words, good reading ability was associated with the expected pattern of reaction times along the stimulus continuum, whereas poor readers were less likely to show the expected pattern; many individual fits showed nearly flat fitted curves or even an inverse pattern to that of the above-average readers. This analysis indicates that children with dyslexia may experience synthesized speech continua in a qualitatively different way from typical readers. Specifically, children with dyslexia are more likely to respond endpoint stimuli and ambiguous stimuli in more or less the same way, whereas above-average readers linger over ambiguous stimuli while categorizing endpoint stimuli more rapidly.

**Figure 4.**
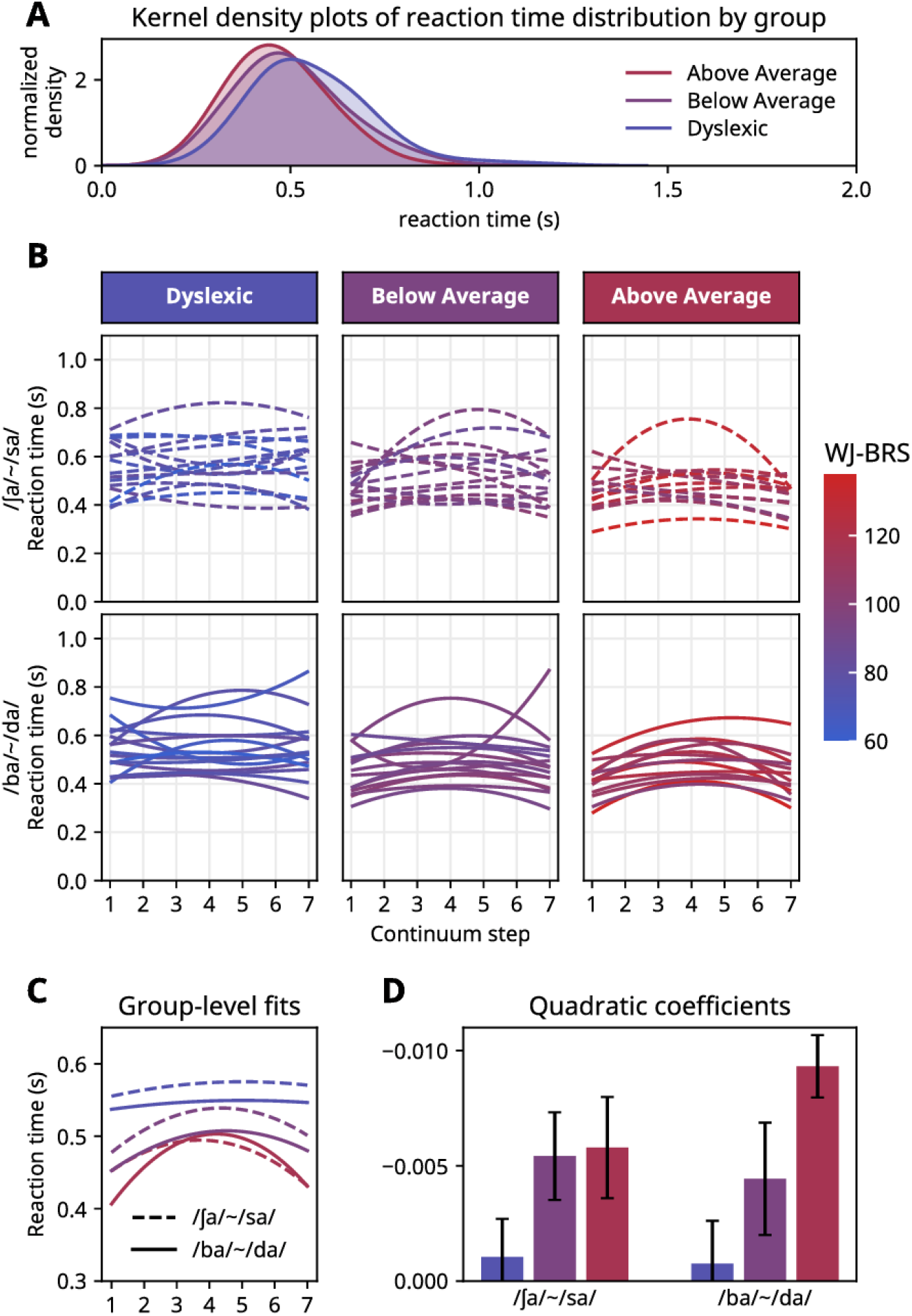
*A: Gaussian kernel density plots (75 ms bandwidth) of raw reaction times pooled across continua and paradigms for each group. B: polynomial fits to reaction times for both speech continua. C: polynomials fit to aggregated participant reaction times for each of the two speech continua. D: mean quadratic coefficients (± 1 standard error of the mean) of the polynomial fits for subjects in each group for the two speech continua*.

### Other measures of reading ability

As a final post-hoc test, we explored which specific components of reading skill might underlie the relationship with performance on speech categorization tasks. To do this, we performed post-hoc Pearson correlation tests between our estimates of psychometric slope, lapse rate, or the principal component that combines slope and asymptote parameters, and several tests targeting specific aspects of reading ability. Those correlations are summarized in Figure 5. The most striking pattern is that the majority of significant correlations between test scores and our model parameters correlate with results from the ABX blocks, but not the single-stimulus blocks. Moreover, it appears that correlations between test scores and lapse rate are clustered on the dynamic-cue /ba/~/da/ continuum, whereas correlations with psychometric slope are clustered on the static-cue /∫a/~/sa/ continuum. Perhaps unsurprisingly, the correlations between test scores and the principal component combining slope and asymptote parameters look something like a union of the correlation patterns for slope and lapse rate. These patterns suggest a striking difference between the ABX paradigm and the single-stimulus paradigm, in how performance on such tasks relates to reading ability.

A second notable feature of these correlations is that tests targeting phonological awareness and phonological memory are no better at predicting children’s psychometric function shapes than tests of real-word or pseudo-word reading. One interpretation of this phenomenon is that, contrary to what has been argued in the literature (Manis *et al.*, 1997; Joanisse *et al.*, 2000; Boets *et al.*, 2010), the phoneme categorization task we used is not in fact a reliable probe of phonological processing (whereas the various test batteries are). If, on the other hand, one accepts that (psychometric models of) children’s performance on phoneme categorization tasks do reflect differences phonological processing, then these results call into question the specificity of the component tests of these various diagnostic batteries.

**Figure 5:**
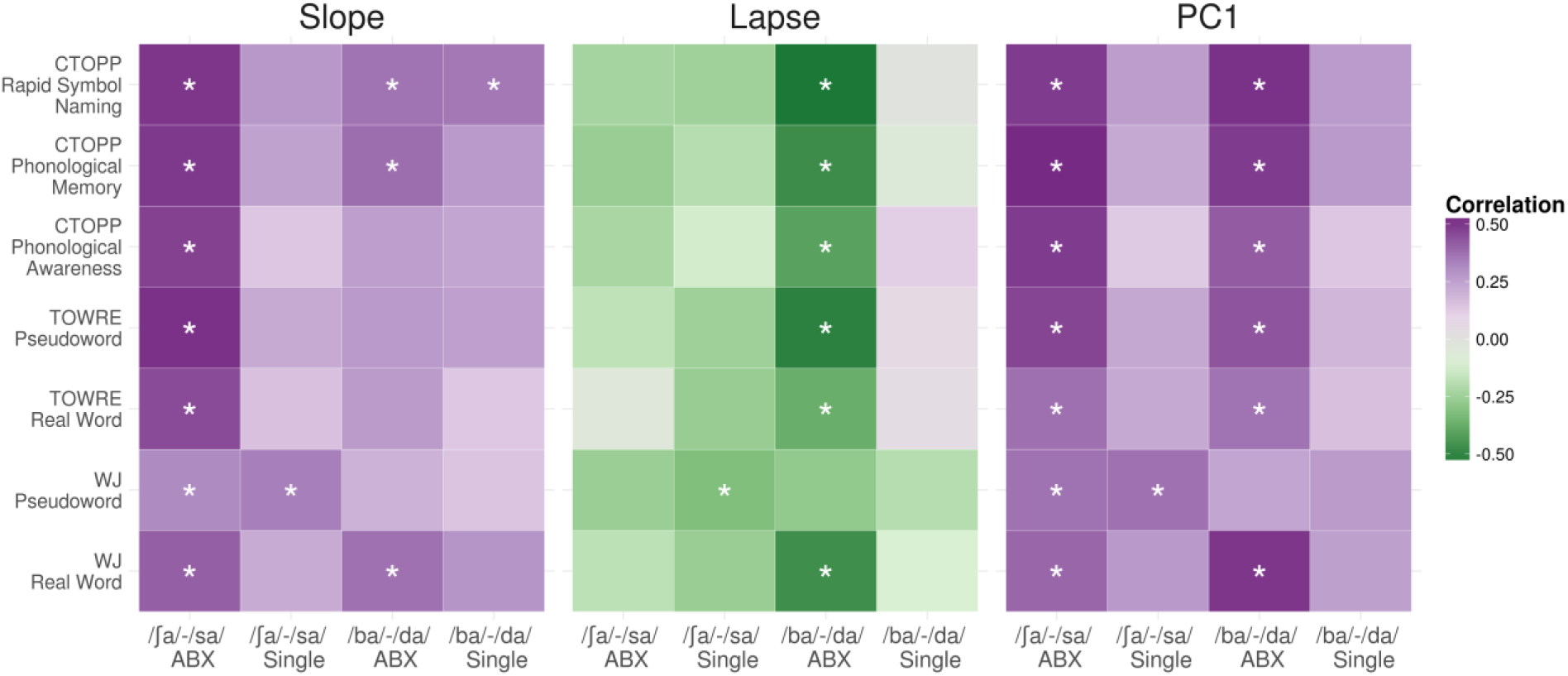
*Pearson correlations between behavioral measures and slope, lapse, and first principal component (PC1) of the psychometric functions estimated from behavioral responses. Color gradient indicates the magnitude of the correlation and asterisks indicate significant correlations (p < 0.05, uncorrected)*.

## Discussion

Many previous studies have claimed to show a relationship between dyslexia and poor categorical perception of speech phonemes (particularly for contrasts that rely on dynamic spectrotemporal cues), while others have suggested that the apparent auditory or linguistic processing impairments are in fact the result of general attention or memory deficits that manifest because of task difficulty. Our data unambiguously show a relationship between reading ability and phoneme categorization performance: higher reading scores were associated with steeper categorization functions, lower lapse rates, faster response times, and greater tendency to respond more quickly for unambiguous stimuli. But we also set out to answer two more specific questions: first, do dyslexic children struggle to categorize sounds on the basis of dynamic auditory cues to a greater extent than static cues? And second, are dyslexic children affected by task difficulty more than typically developing children? We also took care to show the ramifications of methodological choices when modeling categorization data as a psychometric function (particularly the common assumption of zero lapse rate). We discuss each of these points in turn.

### Processing static versus dynamic cues

It has been argued that dyslexics suffer primarily from an auditory processing deficit that specifically affects the processing of temporal information, leading to unique impairments in processing dynamic temporal and spectrotemporal cues such as formant transitions (Vandermosten *et al.*, 2010, 2011) or modulations on the timescale of 2 to 20 Hz (Law *et al.*, 2014). We found that dyslexic children showed, on average, less steep categorization functions regardless of whether they were categorizing on the basis of static or dynamic speech cues. However, estimated lapse rate was related to cue type in our data (though in the opposite direction than what would be predicted from the literature: higher lapse rates were associated with the static cue continuum, not the dynamic one). These results run counter to the findings of Vandermosten and colleagues (2010, 2011), who showed a greater deficit in categorization of speech continua involving dynamic cues compared to static cues. We believe the key difference arises from our choice to equalize the duration of the cues in all stimulus tokens. Vandermosten and colleagues used a 100 ms dynamic cue in their /ba/~/da/ continuum, and a 350 ms static cue in their /i/-/y/ continuum, which raised the possibility that evidence accumulation, not the static or dynamic nature of the speech cue, was source of their observed difference.

One interpretation of our results is that categorizing phonemes may be more difficult when judgments must be made on the basis of very brief cues (whether static or dynamic), and that children with dyslexia are especially susceptible to this difficulty. However, most evidence suggests that slowing down speech does not make it more discriminable for reading and language impaired children (Stollman, Kapteyn and Sleeswijk, 1994; Bradlow *et al.*, 1999). Additionally, we cannot rule out that dyslexics have some impairment specifically in processing syllable-length sounds, as has been recently suggested (Goswami, 2011; Lehongre *et al.*, 2011; Law *et al.*, 2014). What we can conclude is that modulations on the order of 10 Hz (the rate of formant transition changes in our /ba/~/da/ continuum) do not pose a unique difficulty for children with dyslexia, when compared to static speech cues of comparable duration.

### The role of task difficulty

In our data, estimates of psychometric slope and lapse rate were unrelated to whether the task was the more difficult ABX paradigm or the easier single-stimulus paradigm. Additionally, the covariates for ADHD diagnosis and non-verbal IQ were never shown to contribute to better model fits of our data, a contribution we would have expected if task difficulty were a serious issue for our participants. However, our PCA analysis of overall psychometric curve shape provides some evidence that poor readers were more affected by task difficulty than strong readers, and the correlations between various tests of reading ability and psychometric slope or lapse rate were nearly always greater in the ABX paradigm than in the single-stimulus paradigm (see Figure 5).

It is possible that our ABX and single-stimulus conditions were simply not different enough in difficulty to reveal an effect of task difficulty in our main statistical models, despite the apparent difference suggested by the post-hoc correlations with test battery scores. Pertinent to that possibility is a recent meta-analysis of categorical perception studies among children with dyslexia (Noordenbos and Serniclaes, 2015), which found that there is generally a larger effect size reported for discrimination tasks than identification tasks. Indeed, during initial piloting of this study, subjects also performed same/different discrimination task with the same stimulus continua, but performance was so uniformly poor that the discrimination task was dropped from the final experiment. In any case, our findings reinforce claims that in order to fully understand what stimulus-dependent difficulties are faced by people with dyslexia, the stimuli should be tested in more than one context to separate the effects of stimulus properties from the effects of task demands (Banai and Ahissar, 2006).

### The influence of methodological choices

One important point that our study clarifies is the influence of methodological choices on psychometric fitting results. Specifically, we compared 4-parameter models of participants’ psychometric functions with traditional 2-parameter models that ignore lapse rate by fixing the asymptotes at zero and one. Our crossvalidation of the asymptote prior distributions make it clear that a 2-parameter model is a worse fit to the data than any of the possible 4-parameter models we tested (see Materials & Methods, Figure 7). More importantly, using a 2-parameter fit in spite of its poor fit to the data leads to incorrect conclusions: 2-parameter models indicated that slopes were lower for the static cue continuum, whereas 4-parameter models indicated that slope was no different between the two continua, but lapse rate was higher on the static cue continuum. In either case, the main finding—a clear relationship between reading ability and psychometric slope—remains. This is, to a degree, reassuring about previous conclusions drawn from 2-parameter models. Many prior studies may still be correct in finding that children with dyslexia tend to have shallower categorization function slopes on particular types of stimuli. However, by failing to model lapse rate those studies may have overestimated the effect size when comparing children with dyslexia against typical readers.

To further illustrate this point, we explored how the conclusions of our study would have changed if we had changed the assumptions of our psychometric fitting routine. Specifically, we varied the width of the Bayesian priors governing the two asymptotic parameters from 0 (effectively fixed) to 0.5 in steps of 0.05, and observed the effects on the correlation of reading ability and slope. We fit psychometric functions for each subject at each of the eleven prior widths, and at each prior width, we computed the correlations between reading score and slope or lapse rate on the ABX trials. Figure 6 illustrates the result; the apparent correlation between reading ability and psychometric slope is greatest when the priors on the asymptotes are strongest, and the correlation diminishes when the asymptotes are allowed to vary more freely. Thus, it is critical to determine the optimal parameterization of the psychometric function before interpreting the resulting parameters of the fit.

**Figure 6.**
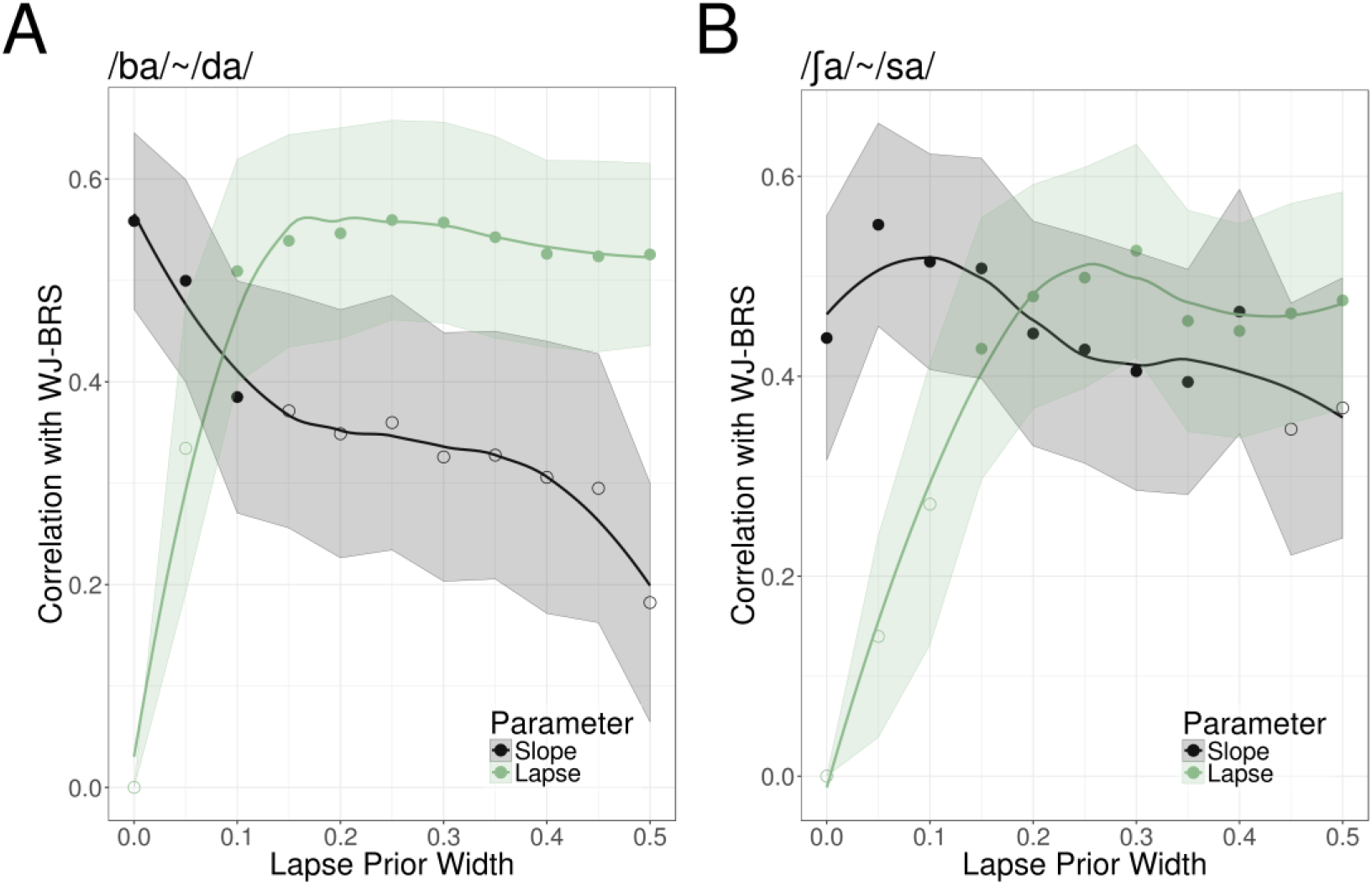
*Correlations between slope, lapse rate, and reading ability (WJ-BRS score) as a function of the asymptotic prior width. Correlations are for psychometric functions fit to ABX trials. Error ribbons are computed by 10,000 bootstrap simulations. Filled dots indicate significant correlations (p < 0.05)*.

### Sensory impairment or language impairment?

What is still not clear, and is not within the scope of this study, is the extent to which low-level sensory abilities drive the relationship between phoneme categorization performance and low reading ability. Gap detection thresholds (the gold standard for measuring temporal resolution) have been shown to be normal in dyslexics (McAnally and Stein, 1997; Adlard and Hazan, 1998; Schulte-Körne *et al.*, 1998; Boets *et al.*, 2007), so poor auditory temporal resolution is not likely to be the underlying cause for the correlation between reading ability and categorization performance. There is also some evidence that backward masking is more severe in dyslexics (Wright *et al.*, 1997), which could potentially mask the consonant cue preceding the vowel in /ba/~/da/ and /∫a/~/sa/ stimuli, and could explain the difference between our findings and those of Vandermosten and colleagues (2010, 2011) regarding static-cue stimuli. However, Rosen and Manganari (2001) showed that differences in backward masking did not affect categorical perception, so there is to date no clear relationship between any measured low-level deficit among poor readers and their ability to categorize speech sounds.

It has been long established that perceptual boundaries typically sharpen as a function of linguistic experience (Garnica, 1973), and categorical perception of speech sounds in pre-lingual children has been shown to predict later success in reading (Bradley and Bryant, 1983). Thus impairment on categorization tasks among children with dyslexia may represent the lower end of a distribution of literacy and categorical perception abilities that occurs in the general population, rather than a specific deficit that is unique to dyslexia (Shaywitz *et al.*, 1992). Our inclusion of children with a large range of reading abilities, rather than just two groups of children widely separated in literacy, made it obvious that there is no clear and distinct boundary between children with dyslexia and “everyone else.” Rather, variation in our models of children’s perceptual boundaries was moderately related to reading ability, and the effect size and variability across our sample argues against interpreting the group-level effect as a unifying feature of dyslexia. Some poor readers had apparently normal psychometric functions, and there was considerable overlap in performance between poor readers with dyslexia and merely below-average readers. That aspect of our findings is in line with a multiple deficit model of dyslexia, whereby multiple underlying mechanisms, including auditory, visual and language deficits, all confer risk for reading difficulties (Pennington, 2006; Peterson and Pennington, 2015; Joo *et al.*, 2017; Joo, Donnelly and Yeatman, 2017).

### Future directions

Our results invite several questions for further exploration. First, although we found that children with dyslexia showed poor performance categorizing speech on the basis of (100 ms) steady-state spectral cues, Vandermosten et al. (2010, 2011) showed that a large sample of dyslexic adults and children were not significantly impaired on a similar task with (350 ms) steady-state vowels. A natural extension of this work would be to parametrically vary cue duration, to look for a duration-dependent difference between readers with and without dyslexia. Such a difference could reflect a difference in sensory integration, which has been probed at the group level in dyslexics for visual stimuli, but not in relation to other sensory modalities (Talcott *et al.*, 2000). Another question raised by our research is how directly auditory phoneme categorization relates to the common diagnostic measure “phonological awareness.” Although it has been hypothesized that poor phoneme categorization reflects fuzzy underlying phonological representations, which in turn mediate reading ability, our study showed only weak correlations between categorization function slope and phonological awareness, compared to the relationship between slope and a compound measure of reading skill. Therefore, it may be worthwhile to perform a more targeted analysis of the relationship between categorical perception of speech and phonological awareness.

## Materials and Methods

All the code for stimulus presentation, data analysis, and figure generation is available at https://github.com/YeatmanLab/Speech_contrasts_public alongside the de-identified data reported in this study.

### Participants

A total of 52 native English-speaking school-aged children with normal hearing were recruited for the study in the study. Children ages 8–12, without histories of neurological or auditory disorders, were recruited from a database of volunteers in the Seattle area (University of Washington Reading & Dyslexia Research Database). Each participant provided written informed consent under a protocol that was approved by the University of Washington Institutional Review Board. All subjects had normal or corrected-to-normal vision. Participants were tested on a battery of assessments, including the Woodcock-Johnson IV (WJ-IV) Letter Word Identification and Word Attack sub-tests, the Test of Word Reading Efficiency (TOWRE), and the Wechsler Abbreviated Scale of Intelligence (WASI-III). All participants underwent a hearing screening to ensure pure tone detection at 500 Hz, 1000 Hz, 2000 Hz, 4000 Hz and 8000 Hz in both ears at 25 dB HL or better. A total of six subjects who were initially recruited did not pass the hearing screening and were not entered into the study, and two more were unable to successfully complete training (described below). Thus a total of 44 children participated in the full experiment.

### Demographics

In order to understand the relationship between phoneme categorization ability and reading ability, we selected our cohort of participants to encompass both impaired and highly skilled readers. Although we treat reading ability as a continuous covariate in our statistical analyses, for the purpose of recruitment and data visualization we defined three groups based on the composite Woodcock-Johnson Basic Reading Score (WJ-BRS) and TOWRE index. The “Dyslexic” group comprised participants who scored 1 standard deviation or more below the mean (standardized score of 100) on both the WJ-BRS and TOWRE index; Above Average readers were defined as those who scored above the mean on both tests; and Below Average readers were defined as participants who fell between the Dyslexia and Above Average groups. We used both the WJ-BRS and TOWRE index in our criterion to improve the confidence of our group assignments, though they are highly correlated measures in our sample (r = 0.89, p <1e−5). There were 15 subjects in the Dyslexic group, 16 in the Below Average group, and 13 in the Above Average group. There were no significant differences in age between groups (Kruskal-Wallis rank sum test, H(2) = 1.0476, p = 0.5923), nor was there a significant correlation between age and WJ-BRS score (r = 0.02, p = 0.89). We did not exclude participants with ADHD diagnoses from the study because ADHD is highly comorbid with dyslexia (Light *et al.*, 1995; Stevenson *et al.*, 2005), so this inclusion contributes to a more representative sample of children. However, we did account for the presence of ADHD diagnosis in our statistical models. Of our 44 participants, 7 had a formal diagnosis of ADHD: 2 in the Above Average group, 1 in the Below Average group, and 4 in the Dyslexic group.

Table 1 shows group comparisons on measures of reading and cognitive skills, and the distributions of each variable are illustrated in Supplementary Figure S1. All subjects had an IQ of at least 80 as measured by the WASI-III FS-2. Therefore, although there was a significant difference in IQ scores across groups, we were not concerned that abnormally low cognitive ability would prevent any child from performing the experimental task. While we expected IQ scores to be higher among stronger readers due to the verbal component of IQ assessment, we also noticed a significant difference in non-verbal IQ across the groups as measured on the WASI-III Matrix Reasoning test. To be certain our results were not confounded by this difference, we also included nonverbal IQ as a covariate in our statistical analyses to confirm the specificity of the relationship with reading skills as opposed to IQ.

**Table 2.**
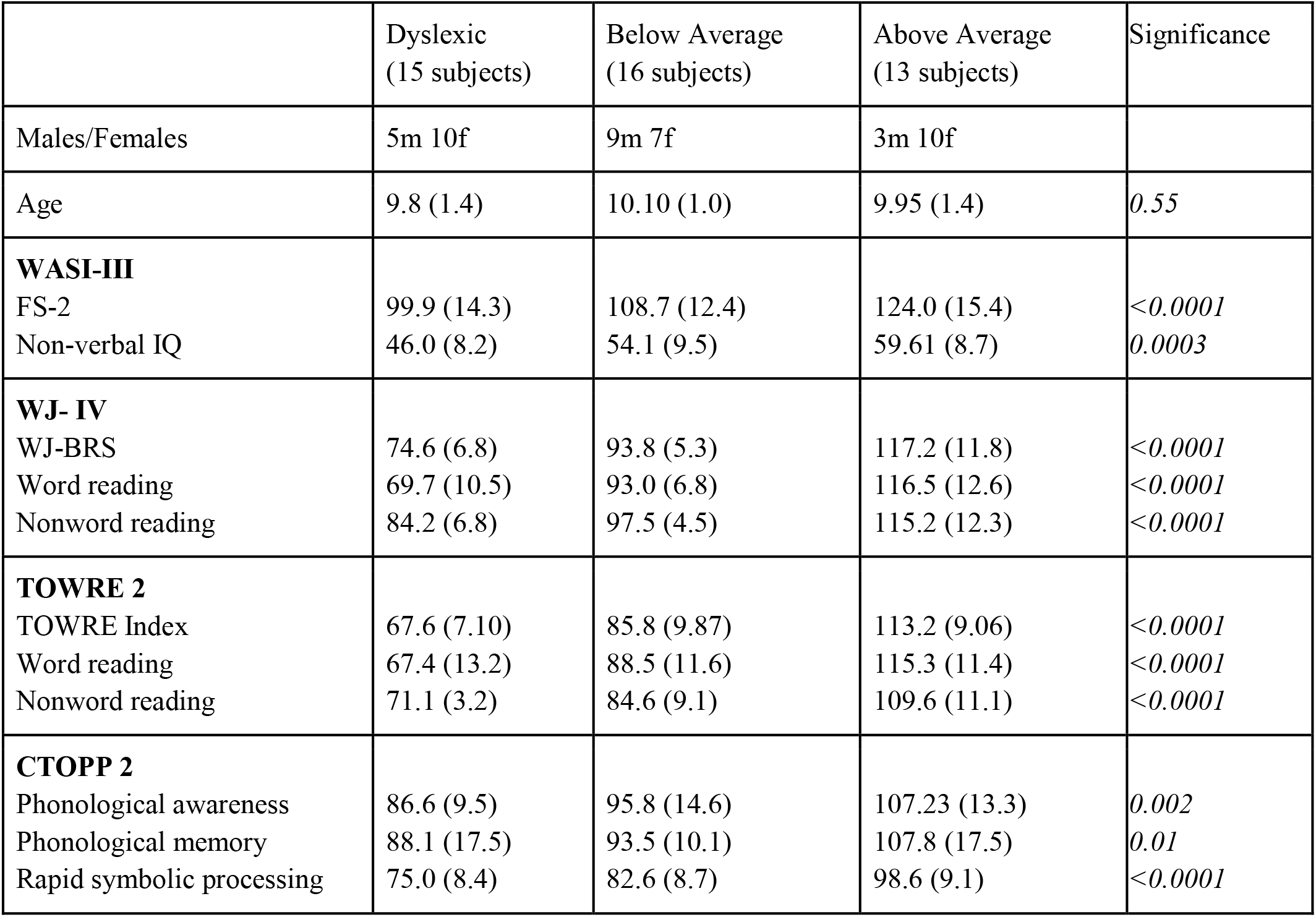
*Summary statistics and group differences on various demographic, reading and cognitive measures. Summary statistics show means and (sample standard deviations). Significance assessed by the Kruskal-Wallis test*.

### Stimuli

Two 7-step speech continua were created using Praat v. 6.0.37 (Boersma, 2001), a /ba/~/da/ continuum and a /∫a/~/sa/ continuum, chosen to probe categorization on the basis of dynamic and static auditory cues, respectively. In the /ba/~/da/ continuum, the starting frequency of the second vowel formant (F2) transition was varied. Individuals who are insensitive to this spectrotemporal modulation should have shallow psychometric functions on the /ba/~/da/ categorization task. The other continuum was /∫a/~/sa/, in which the spectral shape of the fricative noise was varied between tokens (but was static for any given token). Both the /∫a/~/sa/ fricative duration and the /ba/~/da/ F2 transition duration were 100 ms. Endpoint and steady-state formant values for the /ba/~/da/ continuum were measured from recorded syllables from an adult male speaker of American English. The spectral peaks of the /∫a/~/sa/ continuum were chosen from the spectral peaks measured in a recorded /sa/ spoken by the same talker. Spectrograms of the endpoints of both continua are shown in Supplemental Figure S2.

#### The /ba/~/da/ continuum

Synthesis of the /ba/~/da/ continuum followed the procedure described in Winn and Litovsky (2015). Briefly, this procedure involves downsampling a naturally produced /ba/ token, extracting the first four formant contours via linear predictive coding, altering the F2 formant contour to change the starting frequency from 1085 Hz (/ba/) to 1460 Hz (/da/) in seven linearly-spaced steps, linearly interpolating F2 values for the ensuing 100 ms to make a smooth transition to the steady-state portion of the vowel (1225 Hz), and re-generating the speech waveform using Praat’s source-filter synthesis (with a source signal extracted from a neutral vowel from the same talker). Because this procedure eliminates high-frequency energy in the signal at the downsampling step, the synthesized speech sounds are then low-pass filtered at 3500 Hz and combined with a version of the original /ba/ recording that had been high-pass filtered above 3500 Hz. This improves the naturalness of the synthesized sounds, while still ensuring that the only differences between continuum steps are in the F2 formant transitions.

#### The /∫a/~/sa/ continuum

The /∫a/~/sa/continuum was created by splicing synthesized fricatives lasting 100 ms onto a natural /a/ token excised from a spoken /sa/ syllable. The duration of /a/ was scaled to 250 ms using Praat’s implementaion of the PSOLA algorithm (Moulines and Charpentier, 1990). Synthesized fricatives contained three spectral peaks centered at 3000, 6000, and 8000 Hz. The bandwidths and amplitudes of the spectral peaks were linearly interpolated between continuum endpoints in seven steps, and the resulting spectra were used to filter white noise. To improve the naturalness of the synthesized fricatives, a gentle cosine ramp over 75 ms and fall over the final 20 ms was imposed on the fricative envelope. Aside from this onset/offset ramping (which was applied equally to all continuum steps), the contrastive cue (the amplitudes and bandwidths of the spectral peaks) was steady throughout the 100-ms duration of each fricative.

### Procedure

Stimulus presentation and participant response collection was managed with PsychToolbox for MATLAB (Brainard, 1997; The Mathworks Inc., 2016). Auditory stimuli were presented at 75dB SPL via circumaural headphones (Sennheiser HD 600). Children were trained to associate sounds from the two speech continua with animal cartoons on the left and right sides of the screen and to indicate their answers with right or left arrow keypresses. For the /∫a/~/sa/continuum, participants selected between pink and purple snakes. For the /ba/~/da/ continuum, participants selected between two different sheep cartoons. Throughout all blocks, each cartoon was always associated with the same stimulus endpoint. Two experimental conditions were presented: an ABX condition and a single-interval condition.

#### The ABX condition

In ABX blocks, participants performed a two-alternative forced-choice identification task in which they heard three stimuli with a 400 ms ISI and indicated whether the third stimulus (X) was more like the first (A) or second (B) stimulus. Stimuli A and B were always endpoints of the continuum, and during the endpoint presentations, the animals associated with those sounds lit up. Based on pilot data with adult participants, varying endpoint order trial-by-trial (i.e., interleaving /ba/-/da/-X and /da/-/ba/-X trials in the same block) was deemed very difficult, so for our young listeners we fixed the order of endpoint stimulus presentation within each block. The order of blocks was counterbalanced across participants. Each block contained 10 trials for each of the seven continuum steps, for a total of 70 trials per block.

#### The single interval condition

In single interval blocks, participants heard a single syllable and decided which category it belonged to by selecting an animal. No text labels were used, so participants learned to associate each animal with a continuum endpoint during a practice round. Two subjects (neither in the Dyslexia group) lost track of the animals associated with the endpoints after one successful practice round and were given brief additional instruction before successfully completing the task. Each block contained 10 trials for each step on the continuum, for a total of 70 trials.

There were thus six blocks total: an ABX /ba/~/da/ block, an ABX /∫a/~/sa/ block, a second ABX /ba/~/da/ block, a second ABX /∫a/~/sa/ block, a single-interval /ba/~/da/ block, and finally a single-interval /∫a/~/sa/ block. Other than the counterbalancing of ABX endpoint order mentioned above, the order of blocks was the same for all participants. Practice rounds were administered before the first ABX round and before each of the single-interval blocks. In practice rounds, participants were asked to categorize only endpoint stimuli and were given feedback on every trial. Participants had to score at least 75% correct on the practice round to advance to the experiment and were allowed to repeat the practice blocks up to three times. Two children did not meet this criterion and were not included in the study.

### Psychometric Curve fitting

Modeling of response data was performed with Psignifit 4.0 (Schütt *et al.*, 2015), a MATLAB toolbox that implements Bayesian inference to fit psychometric functions. We fit a logistic curve with four parameters, modeling the two asymptotes, the width of the psychometric, and the threshold. The width parameter was later transformed to the slope at the threshold value. Four-parameter fits of each experimental block were performed with each of 11 possible priors: a prior that fixed the lapse rate at zero (in line with the approach in most previous studies of auditory processing in dyslexia), and ten uniform distribution priors with lower bound zero and upper bound ranging from 5% to 50% in steps of 5% (e.g., a lapse rate of 20% corresponds to constraining the lower asymptote between 0 and 0.2, and the upper asymptote between 0.8 and 1). Next, the optimal prior was chosen using leave-one-out cross-validation. Specifically, for each of the 261 test blocks across all participants, psychometric curves were fit to 69 of the 70 data points in that block, and the likelihood of the participant’s selection on the held-out data point was calculated under the model. This process was repeated for each of the 70 data points in each block, for each of the 11 possible prior widths. The estimated likelihoods of the held-out data points were used as a goodness-of-fit metric and pooled across blocks and cross-validation runs to determine the median likelihood for each prior width. The optimal prior was determined to be a maximum lapse rate of 10%, as seen in Figure 7; note that when the asymptotes were fixed at 0 and 1, the psychometric fits had the poorest fit to the data.

**Figure 7.**
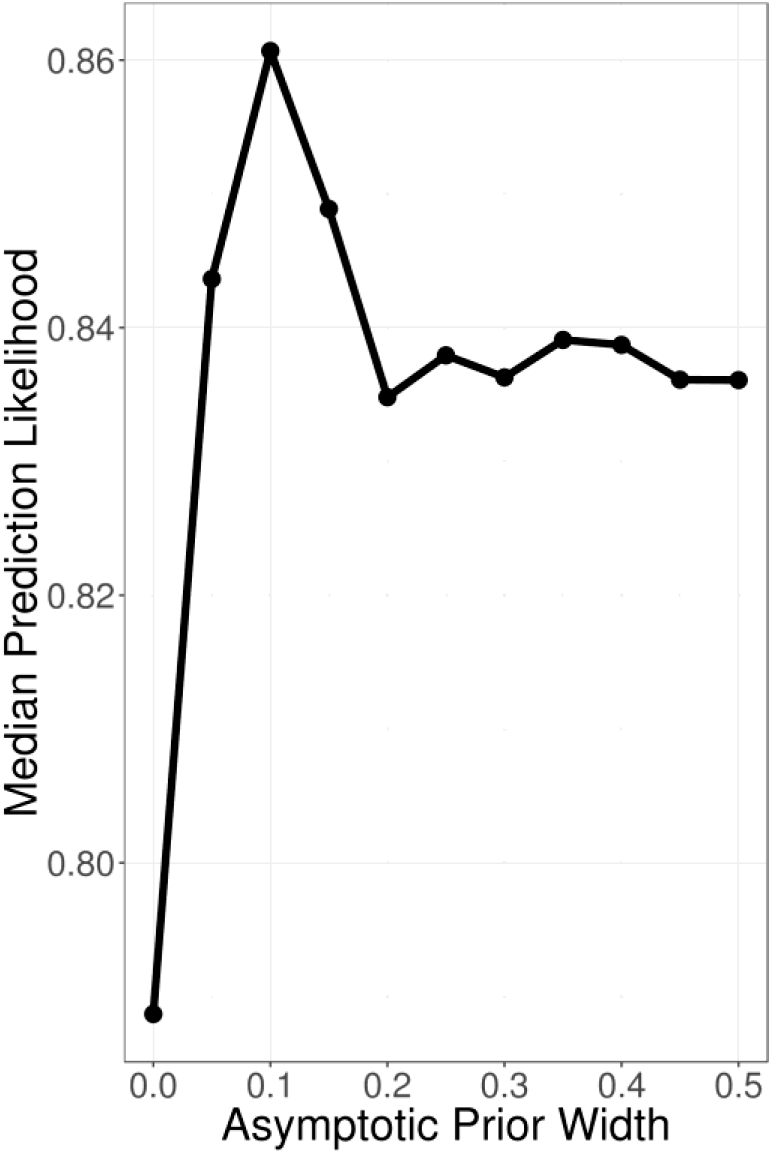
*The median likelihood of held-out data points from all cross-validation trials is plotted against the width of the asymptotic parameter prior distributions (leave-one-out cross validation). The optimal prior width is clearly shown to be one that allows lapse rate to vary between 0 and 0.1. However, in all cases the 4-parameter model fit the data better than the 2-parameter model*.

Having determined the optimal prior for the asymptote parameters, models were re-fit to obtain final estimates of the four parameters, for each combination of participant, cue continuum, and test paradigm. Additionally, 2-parameter fits (with asymptotes fixed at 0 and 1) were also recomputed, for later comparison with the optimal 4-parameter models. Any cases where the best-fit threshold parameter was not in the range of the stimulus continuum steps (1 through 7) were excluded from further analysis. Of the 261 total psychometric functions fit in the study, 12 of the 2-parameter fits and 10 of the 4-parameter fits were excluded on these grounds.

### Reaction time analysis

Post-hoc analysis of reaction time data was also performed. To analyze reaction time, we first removed any trials with a reaction time shorter than 200 ms or longer than 2 s. The lower cutoff was based on the minimum response time to an auditory stimulus (Fry, 1975) such responses are assumed to be spurious or accidental button presses, and resulted in the exclusion of ~9% of trials. The upper cutoff was chosen based on the histograms of reaction times to exclude trials where subjects became distracted and were clearly temporarily disengaged from the experiment, and resulted in the exclusion of ~6% of trials. We fit each individual’s remaining data as a function of continuum step with a 2nd order orthogonal polynomial basis (see Results for rationale), reading ability, and stimulus continuum with a mixed-effects regression model. For visualization, the reaction time data from each experimental block for each participant were reduced by fitting a χ^2^ distribution to the reaction times at each continuum step, and using the peak of the fitted distribution as our summary statistic (because reaction time distributions tend to be right-skewed, making the mean a poor measure of central tendency).

## Acknowledgements

We would like to thank Richard Wright for helpful discussions and feedback in the study design and interpretation. This work was supported by NIH grant 2T32DC005361-16 and research grants from Microsoft to J.D.Y.

## References

Adlard A. and Hazan V. (1998) ‘Speech Perception in Children With Specific Reading Difficulties (Dyslexia)’, The Quarterly Journal of Experimental Psychology Section A, 51(1), pp. 153–177. doi: 10.1080/713755750.

Ahissar M. et al. (2006) ‘Dyslexia and the failure to form a perceptual anchor’, Nature Neuroscience, 9(12), pp. 1558–1564. doi: 10.1038/nn1800.

Ahissar M. (2007) ‘Dyslexia and the anchoring-deficit hypothesis’, Trends in Cognitive Sciences, 11(11), pp. 458–465. doi: 10.1016/j.tics.2007.08.015.

Amitay S. et al. (2002) ‘Disabled readers suffer from visual and auditory impairments but not from a specific magnocellular deficit.’, Brain: a journal of neurology, 125(Pt 10), pp. 2272–2285. doi: 10.1093/brain/awf231.

Banai K. and Ahissar M. (2004) ‘Poor frequency discrimination probes dyslexics with particularly impaired working memory’, Audiology and Neuro-Otology, 9(6), pp. 328–340. doi: 10.1159/000081282.

Banai K. and Ahissar M. (2006) ‘Auditory processing deficits in dyslexia: Task or stimulus related?’, Cerebral Cortex, 16(12), pp. 1718–1728. doi: 10.1093/cercor/bhj107.

Bates D. et al. (2015) ‘Fitting Linear Mixed-Effects Models using lme4’, Journal of Statistical Software, 67(1), pp. 1–48. doi: 10.18637/jss.v067.i01.

Boersma P. (2001) ‘Praat, a system for doing phonetics by computer’, Glot International, 5(9/10), pp. 341–347. doi: 10.1097/AUD.0b013e31821473f7.

Boets B. et al. (2007) ‘Auditory processing, speech perception and phonological ability in pre-school children at high-risk for dyslexia: A longitudinal study of the auditory temporal processing theory’, Neuropsychologia, 45(8), pp. 1608–1620. doi: 10.1016/j.neuropsychologia.2007.01.009.

Boets B. et al. (2010) ‘Towards a further characterization of phonological and literacy problems in Dutch-speaking children with dyslexia’, British Journal of Developmental Psychology, 28(1), pp. 5–31. doi: 10.1348/026151010X485223.

Boets B. et al. (2011) ‘Preschool impairments in auditory processing and speech perception uniquely predict future reading problems’, Research in Developmental Disabilities, 32(2), pp. 560–570. doi: 10.1016/j.ridd.2010.12.020.

Bogliotti C. et al. (2008) ‘Discrimination of speech sounds by children with dyslexia: Comparisons with chronological age and reading level controls’, Journal of Experimental Child Psychology, 101(2), pp. 137–155. doi: 10.1016/j.jecp.2008.03.006.

Bradley L. and Bryant P. E. (1983) ‘Categorizing sounds and learning to read - A causal connection’, Nature, 301(5899), pp. 419–421. doi: 10.1038/301419a0.

Bradlow A. R. et al. (1999) ‘Effects of lengthened formant transition duration on discrimination and neural representation of synthetic CV syllables by normal and learning-disabled children’, The Journal of the Acoustical Society of America, 106(4), p. 2086. doi: 10.1121/1.427953.

Brainard D. H. (1997) ‘The Psychophysics Toolbox’, Spatial Vision, 10(4), pp. 433–436. doi: 10.1163/156856897X00357.

Breier J. I. et al. (2001) ‘Perception of Voice and Tone Onset Time Continua in Children with Dyslexia with and without Attention Deficit/Hyperactivity Disorder’, Journal of Experimental Child Psychology, 80(3), pp. 245–270. doi: 10.1006/jecp.2001.2630.

Bus A. G. and van IJzendoorn M. H. (1999) ‘Phonological awareness and early reading: A meta-analysis of experimental training studies.’, Journal of Educational Psychology, 91(3), pp. 403–414. doi: 10.1037//0022-0663.91.3.403.

Chiappe, P., Chiappe, D. L. and Siegel, L. S. (2001) ‘Speech Perception, Lexicality, and Reading Skill’, Journal of Experimental Child Psychology, 80(1), pp. 58–74. doi: 10.1006/jecp.2000.2624.

Dawes P. et al. (2009) ‘Temporal auditory and visual motion processing of children diagnosed with auditory processing disorder and dyslexia’, Ear and Hearing, 30(6), pp. 675–686. doi: 10.1097/AUD.0b013e3181b34cc5.

Farmer M. E. and Klein R. M. (1995) ‘The evidence for a temporal processing deficit linked to dyslexia: A review’, Psychonomic Bulletin & Review, 2(4), pp. 460–493. doi: 10.3758/BF03210983.

Fry D. B. (1975) ‘Simple Reaction-Times to Speech and Non-Speech Stimuli’, Cortex, 11(4), pp. 355–360. doi: 10.1016/S0010-9452(75)80027-X.

Garnica O. K. (1973) ‘The development of phonemic speech perception’, in Cognitive development and the acquisition of language, pp. 215–222.

Gibson, L. Y., Hogben, J. H. and Fletcher, J. (2006) ‘Visual and auditory processing and component reading skills in developmental dyslexia’, Cognitive Neuropsychology, 23(4), pp. 621–642. doi: 10.1080/02643290500412545.

Goswami U. et al. (2002) ‘Amplitude envelope onsets and developmental dyslexia: A new hypothesis’, Proceedings of the National Academy of Sciences, 99(16), pp. 10911–10916. doi: 10.1073/pnas.122368599.

Goswami U. (2011) ‘A temporal sampling framework for developmental dyslexia’, Trends in Cognitive Sciences. Elsevier Ltd, 15(1), pp. 3–10. doi: 10.1016/j.tics.2010.10.001.

Goswami U. et al. (2011) ‘Rise time and formant transition duration in the discrimination of speech sounds: The Ba-Wa distinction in developmental dyslexia’, Developmental Science, 14(1), pp. 34–43. doi: 10.1111/j.1467-7687.2010.00955.x.

Hämäläinen J. A. et al. (2009) ‘Common variance in amplitude envelope perception tasks and their impact on phoneme duration perception and reading and spelling in Finnish children with reading disabilities’, Applied Psycholinguistics, 30(3), pp. 511–530. doi: 10.1017/S0142716409090250.

Hämäläinen, J. A., Salminen, H. K. and Leppänen, P. H. T. (2013) ‘Basic Auditory Processing Deficits in Dyslexia’, Journal of Learning Disabilities, 46(5), pp. 413–427. doi: 10.1177/0022219411436213.

Hulme C. et al. (2002) ‘Phoneme awareness is a better predictor of early reading skill than onset-rime awareness’, Journal of Experimental Child Psychology, 82(1), pp. 2–28. doi: 10.1006/jecp.2002.2670.

Van Ingelghem, Mieke, Boets, B. et al. (2005) ‘An auditory temporal processing deficit in children with dyslexia’, in Learning disabilities: A challenge to teaching and instruction. Series: Studia Paedagogica, pp. 47–63.

Joanisse M. F. et al. (2000) ‘Language Deficits in Dyslexic Children: Speech Perception, Phonology, and Morphology’, Journal of Experimental Child Psychology, 77(1), pp. 30–60. doi: 10.1006/jecp.1999.2553.

Joo S. J. et al. (2017) ‘Optimizing text for an individual’s visual system: The contribution of crowding to reading difficulties’, CORTEX. Elsevier Ltd, (April), pp. 1–27. doi: 10.1016/j.cortex.2018.03.013.

Joo, S. J., Donnelly, P. M. and Yeatman, J. D. (2017) ‘The causal relationship between dyslexia and motion perception reconsidered’, Scientific Reports, 7(1). doi: 10.1038/s41598-017-04471-5.

Law J. M. et al. (2014) ‘The relationship of phonological ability, speech perception, and auditory perception in adults with dyslexia’, Frontiers in Human Neuroscience, 8. doi: 10.3389/fnhum.2014.00482.

Lehongre K. et al. (2011) ‘Altered low-gamma sampling in auditory cortex accounts for the three main facets of dyslexia’, Neuron, 72(6), pp. 1080–1090. doi: 10.1016/j.neuron.2011.11.002.

Light J. G. et al. (1995) ‘Reading disability and hyperactivity disorder: Evidence for a common genetic etiology’, Developmental Neuropsychology, 11(3), pp. 323–335.

Lyon, G. R., Shaywitz, S. E. and Shaywitz, B. A. (2003) ‘A definition of dyslexia’, Annals of Dyslexia, 53(1), pp. 1–14. doi: 10.1007/s11881-003-0001-9.

Maassen B. et al. (2001) ‘Identification and discrimination of voicing and place-of-articulation in developmental dyslexia’, Clinical Linguistics and Phonetics, 15(4), pp. 319–339. doi: 10.1080/02699200010026102.

Manis F. R. et al. (1997) ‘Are Speech Perception Deficits Associated with Developmental Dyslexia?’, Journal of Experimental Child Psychology, 66(2), pp. 211–235. doi: 10.1006/jecp.1997.2383.

McAnally K. I. and Stein J. F. (1997) ‘Scalp potentials evoked by amplitude-modulated tones in dyslexia.’, Journal of speech, language, and hearing research: JSLHR, 40, pp. 939–945. doi: 10.1044/jslhr.4004.939.

Menell, P., McAnally, K. I. and Stein, J. F. (1999) ‘Psychophysical sensitivity and physiological response to amplitude modulation in adult dyslexic listeners’, Journal of Speech, Language, and Hearing …, 42(4), pp. 797–803. doi: 10.1044/jslhr.4204.797.

Merzenich M. M. et al. (1996) ‘Temporal processing deficits of language-learning impaired children ameliorated by training.’, Science (New York, N.Y.), 271(5245), pp. 77–81. doi: 10.1126/science.271.5245.77.

Messaoud-Galusi, S., Hazan, V. and Rosen, S. (2011) ‘Investigating Speech Perception in Children With Dyslexia: Is There Evidence of a Consistent Deficit in Individuals?’, Journal of Speech Language and Hearing Research, 54(6), p. 1682. doi: 10.1044/1092-4388(2011/09-0261).

Moulines, É. and Charpentier, F. (1990) ‘Pitch-synchronous waveform processing techniques for text-to-speech synthesis using diphones’, Speech Communication, 9(5-6), pp. 453–467. doi: 10.1016/0167-6393(90)90021-Z.

Noordenbos M. W. and Serniclaes W. (2015) ‘The Categorical Perception Deficit in Dyslexia: A MetaAnalysis’, Scientific Studies of Reading, 19(5), pp. 340–359. doi: 10.1080/10888438.2015.1052455.

Pennington B. F. (2006) ‘From single to multiple deficit models of developmental disorders’, Cognition, 101(2), pp. 385–413. doi: 10.1016/j.cognition.2006.04.008.

Peterson R. L. and Pennington B. F. (2015) ‘Developmental Dyslexia’, Annual Review of Clinical Psychology, 11(1), pp. 283–307. doi: 10.1146/annurev-clinpsy-032814-112842.

Pisoni D. B. and Tash J. (1974) ‘Reaction times to comparisons within and across phonetic categories’, Perception & Psychophysics, 15(2), pp. 285–290. doi: 10.3758/BF03213946.

Poelmans H. et al. (2011) ‘Reduced sensitivity to slow-rate dynamic auditory information in children with dyslexia’, Research in Developmental Disabilities, 32(6), pp. 2810–2819. doi: 10.1016/j.ridd.2011.05.025.

Ramus F. et al. (2003) ‘Theories of developmental dyslexia: Insights from a multiple case study of dyslexic adults’, Brain, 126(4), pp. 841–865. doi: 10.1093/brain/awg076.

Ramus F. and Szenkovits G. (2008) ‘What phonological deficit?’, in Quarterly Journal of Experimental Psychology, pp. 129–141. doi: 10.1080/17470210701508822.

Reed M. A. (1989) ‘Speech perception and the discrimination of brief auditory cues in reading disabled children’, Journal of Experimental Child Psychology, 48(2), pp. 270–292. doi: 10.1016/0022-0965(89)90006-4.

Repp B. H. (1981) ‘Perceptual equivalence of two kinds of ambiguous speech stimuli’, Bulletin of the Psychonomic Society, 18(1), pp. 12–14. doi: 10.3758/BF03333556.

Roach, N. W., Edwards, V. T. and Hogben, J. H. (2004) ‘The tale is in the tail: An alternative hypothesis for psychophysical performance variability in dyslexia’, Perception, 33(7), pp. 817–830. doi: 10.1068/p5207.

Rocheron I. et al. (2002) ‘Temporal envelope perception in dyslexic children.’, Neuroreport, 13(13), pp. 1683–7. doi: 10.1097/00001756-200209160-00023.

Rosen S. (1999) ‘Language disorders: A problem with auditory processing?’, Current Biology, 9(18). doi: 10.1016/S0960-9822(99)80443-6.

Rosen S. (2003) ‘Auditory processing in dyslexia and specific language impairment: Is there a deficit? What is its nature? Does it explain anything?’, Journal of Phonetics, pp. 509–527. doi: 10.1016/S0095-4470(03)00046-9.

Rosen S. and Mangari E. (2001) ‘Speech and Nonspeech Auditory Processing in Children With Dyslexia ?’, Journal of speech, language and hearing research, 44(August), pp. 720–736.

Schulte-Körne G. et al. (1998) ‘Role of auditory temporal processing for reading and spelling disability’, Perceptual and motor skills, 86(3 Pt 1), pp. 1043–1047. doi: 10.2466/pms.1998.86.3.1043.

Schütt H. et al. (2015) ‘Psignifit 4: Pain-free Bayesian Inference for Psychometric Functions.’, Journal of vision, 15(12), p. 474. doi: 10.1167/15.12.474.

Shaywitz S. E. et al. (1992) ‘Evidence That Dyslexia May Represent the Lower Tail of a Normal Distribution of Reading Ability’, New England Journal of Medicine, 326(3), pp. 145–150. doi: 10.1056/NEJM199201163260301.

Shaywitz S. E. (1998) ‘Dyslexia.’, The New England journal of medicine, 338(5), pp. 307–312. doi: 10.1056/NEJM199801293380507.

Siegel L. S. and Ryan E. B. (1989) ‘Subtypes of developmental dyslexia: The influence of definitional variables’, Reading and Writing, 1(3), pp. 257–287. doi: 10.1007/BF00377646.

Snowling M. J. (1998) ‘Dyslexia as a Phonological Deficit: Evidence and Implications’, Child Psychology and Psychiatry Review, 3(1), pp. 4–11. doi: 10.1017/S1360641797001366.

Snowling M. J. (2000) Dyslexia, 2nd Edition, Wiley-Blackwell. Available at: http://www.wiley.com/WileyCDA/WileyTitle/productCd-0631205748.html.

Steinbrink C. et al. (2014) ‘Development of rapid temporal processing and its impact on literacy skills in primary school children’, Child Development, 85(4), pp. 1711–1726. doi: 10.1111/cdev.12208.

Stevenson J. et al. (2005) ‘Attention deficit hyperactivity disorder with reading disabilities: Preliminary genetic findings on the involvement of the ADRA2A gene’, Journal of Child Psychology and Psychiatry and Allied Disciplines, 46(10), pp. 1081–1088. doi: 10.1111/j.1469-7610.2005.01533.x.

Stollman, M. H. P., Kapteyn, T. S. and Sleeswijk, B. W. (1994) ‘Effect of time-scale modification of speech on the speech recognition threshold in noise for hearing-impaired and language-impaired children’, Scandinavian Audiology, 23(1), pp. 39–46. doi: 10.3109/01050399409047484.

Stoodley C. J. et al. (2006) ‘Auditory event-related potentials differ in dyslexics even when auditory psychophysical performance is normal’, Brain Research, 1121(1), pp. 190–199. doi: 10.1016/j.brainres.2006.08.095.

Swanson H. L. (1993) ‘Working memory in learning disability subgroups’, Journal of Experimental Child Psychology, 56(1), pp. 87–114. doi: 10.1006/jecp.1993.1027.

Talcott J. B. et al. (2000) ‘Visual motion sensitivity in dyslexia: Evidence for temporal and energy integration deficits’, Neuropsychologia, 38(7), pp. 935–943. doi: 10.1016/S0028-3932(00)00020-8.

Tallal P. (1980) ‘Auditory temporal perception, phonics, and reading disabilities in children’, Brain and Language, 9(2), pp. 182–198. doi: 10.1016/0093-934X(80)90139-X.

Tallal P. et al. (1996) ‘Language comprehension in language-learning impaired children improved with acoustically modified speech.’, Science, 271(5245), pp. 81–84. doi: 10.1126/science.271.5245.81.

The Mathworks Inc. (2016) MATLAB-MathWorks, www.mathworks.com/products/matlab. doi: 2016-11-26.

Thomson J. M. et al. (2006) ‘Auditory and motor rhythm awareness in adults with dyslexia’, Journal of Research in Reading, 29(3), pp. 334–348. doi: 10.1111/j.1467-9817.2006.00312.x.

Thomson J. M. and Goswami U. (2008) ‘Rhythmic processing in children with developmental dyslexia:Auditory and motor rhythms link to reading and spelling’, Journal of Physiology Paris, 102(1-3), pp. 120–129. doi: 10.1016/j.jphysparis.2008.03.007.

Treutwein B. and Strasburger H. (1999) ‘Fitting the psychometric function’, Perception and Psychophysics, 61(1), pp. 87–106. doi: 10.3758/BF03211951.

Vandermosten M. et al. (2010) ‘Adults with dyslexia are impaired in categorizing speech and nonspeech sounds on the basis of temporal cues’, Proceedings of the National Academy of Sciences, 107(23), pp. 10389–10394. doi: 10.1073/pnas.0912858107.

Vandermosten M. et al. (2011) ‘Impairments in speech and nonspeech sound categorization in children with dyslexia are driven by temporal processing difficulties’, Research in Developmental Disabilities, 32(2), pp. 593–603. doi: 10.1016/j.ridd.2010.12.015.

Vargo, F. E., Grosser, G. S. and Spafford, C. S. (1995) ‘Digit span and other WISC-R scores in the diagnosis of dyslexia in children.’, Perceptual and motor skills, 80(3 Pt 2), pp. 1219–29. doi: 10.2466/pms.1995.80.3c.1219.

Wang S. and Gathercole S. E. (2013) ‘Working memory deficits in children with reading difficulties: Memory span and dual task coordination’, Journal of Experimental Child Psychology, 115(1), pp. 188–197. doi: 10.1016/j.jecp.2012.11.015.

Wichmann F. A. and Hill N. J. (2001a) ‘The psychometric function: I. Fitting, sampling, and goodness of fit’, Perception & Psychophysics, 63(8), pp. 1293–1313. doi: 10.3758/BF03194544.

Wichmann F. A. and Hill N. J. (2001b) ‘The psychometric function: II. Bootstrap-based confidence intervals and sampling’, Perception & Psychophysics, 63(8), pp. 1314–1329. doi: 10.3758/BF03194545.

Winn M. B. and Litovsky R. Y. (2015) ‘Using speech sounds to test functional spectral resolution in listeners with cochlear implants’, The Journal of the Acoustical Society of America, 137(3), pp. 1430–1442. doi: 10.1121/1.4908308.

Witton C. (1998) ‘Sensitivity to dynamic auditory and visual stimuli predicts nonword reading ability in both dyslexic and normal readers’, Current Biology, 8(14), pp. 791–797. doi: 10.1016/S0960-9822(98)70320-3.

Witton C. et al. (2002) ‘Separate influences of acoustic AM and FM sensitivity on the phonological decoding skills of impaired and normal readers.’, Journal of cognitive neuroscience, 14(6), pp. 866–874. doi: 10.1162/089892902760191090.

Wright B. A. et al. (1997) ‘Deficits in auditory temporal and spectral resolution in language-impaired children’, Nature, 387(6629), pp. 176–178. doi: 10.1038/387176a0.

Zhang Y. et al. (2012) ‘Universality of categorical perception deficit in developmental dyslexia: An investigation of Mandarin Chinese tones’, Journal of Child Psychology and Psychiatry and Allied Disciplines, 53(8), pp. 874–882. doi: 10.1111/j.1469-7610.2012.02528.x.

Ziegler J. C. (2008) ‘Better to lose the anchor than the whole ship’, Trends in Cognitive Sciences, pp. 244–245. doi: 10.1016/j.tics.2008.04.001.

